# Diverse modes of binocular interactions in the mouse superior colliculus

**DOI:** 10.1101/2020.12.14.422574

**Authors:** Ashley L. Russell, Karen G. Dixon, Jason W. Triplett

## Abstract

The superior colliculus (SC) integrates visual and other sensory information to regulate critical reflexive and innate behaviors, such as prey capture. In the mouse, the vast majority of retinal ganglion cells (RGCs) innervate the SC, including inputs from both the contralateral (contra-RGCs) and ipsilateral (ipsi-RGCs) eye. Despite this, previous studies revealed minimal neuronal responses to ipsilateral stimulation and few binocular interactions in the mouse SC. More recent work suggests that ipsi-RGC function and innervation of the SC are critical for efficient prey capture, raising the possibility that binocular interactions in the mouse SC may be more prevalent than previously thought. To explore this possibility, we investigated eye-specific and binocular influences on visual responses and tuning of SC neurons, focusing on the anteromedial region. While the majority of SC neurons were primarily driven by contralateral eye stimulation, we observed that a substantial proportion of units were influenced or driven by ipsilateral stimulation. Clustering based on differential responses to eye-specific stimulus presentation revealed five distinct putative subpopulations and multiple modes of binocular interaction, including facilitation, summation, and suppression. Each of the putative subpopulations exhibited selectivity for orientation, and differences in spatial frequency tuning and spatial summation properties were observed between subpopulations. Further analysis of orientation tuning under different ocular conditions supported differential modes of binocular interaction between putative subtypes. Taken together, these data suggest that binocular interactions in the mouse SC may be more prevalent and diverse than previously understood.

## Introduction

The mouse superior colliculus (SC) has emerged as an attractive model to interrogate visual circuits from developmental, functional, and behavioral perspectives (1, 2). The mouse SC is a primary target of retinal innervation, receiving projections from 85-90% of retinal ganglion cells (RGCs) (3). The vast majority of these inputs originate from the contralateral eye (contra-RGCs), but RGCs from the ipsilateral eye (ipsi-RGCs), which represent only ^~^5% of all RGCs (4), also innervate the SC. Within the superficial layers of the SC, ipsi-RGCs terminate in a sublayer beneath contra-RGCs and are topographically localized to a patchy crescent along the anteromedial border (5). Despite being innervated by both contra- and ipsi-RGCs, little is known about how eye-specific inputs influence visual processing in the rodent SC. Indeed, early studies in the mouse (6) and hamster (7) reported little influence of visual responses by ipsilateral stimulation. However, concurrent studies in the hamster SC in which the anteromedial region was specifically sampled revealed a greater proportion of ipsilaterally-driven neurons (8). But, outside of the ability to drive SC neuron firing, little other characterization of binocularly-modulated visual responses was reported.

The organization of eye-specific inputs into segregated layers and the patchy innervation patterns of ipsi-RGCs might suggest minimal crossover in the influence of each eye on visual function in the SC. However, recent studies suggest that binocular interactions within the SC may play an important role in the critical innate behavior of prey capture. To begin, prey capture is dependent on vision (9), and distinct subtypes of SC neurons regulate different aspects of the behavior (10). Intriguingly, prey capture requires binocular vision and the function of ipsilaterally-projecting retinal ganglion cells (ipsi-RGCs) (11). Furthermore, ipsi-RGC projections to the SC are required for efficient prey capture (12), raising the possibility that binocular interactions within the SC play a critical role in this behavior. However, we know surprisingly little about the ways in which eye-specific inputs modulate visual function in the mouse SC.

Here, we utilized *in vivo* electrophysiological recordings in anesthetized mice to characterize the responses of neurons in the anteromedial portion of the SC to visual stimuli presented to each eye individually or both together. Consistent with previous studies, we found that the majority of neurons were predominantly driven by the contralateral eye. However, a subset of these were facilitated in binocular viewing conditions. Furthermore, we observed a substantial population of neurons which were driven to a great degree by ipsilateral stimulation. Of these, one putative subtype exhibited additive responses under binocular viewing conditions, another exhibited robust facilitation, and a third exhibited suppression by the contralateral stimulation. Across the five putative subtypes of ocular modulation, we observed robust tuning to orientation, but little selectivity for direction of movement. Further analyses of orientation-selective neurons revealed differences in spatial frequency tuning and linearity of response between putative subtypes. Finally, analysis of orientation tuning under distinct ocular viewing conditions revealed potential modes of binocular interactions in each putative subtype. Taken together, these data suggest that binocular interactions in the mouse SC may be more prevalent and varied than previously understood.

## Materials and Methods

### Ethical approval

All the experimental procedures, care and handling of animals were conducted in accordance with and approved by the Institutional Animal Care and Use Committee at Children’s National Hospital.

### Subjects

Adult (45-120 days of age) male and female C57BL/6J mice (RRID: IMSR_JAX:000664, Jackson Laboratory) were used for this study. Mice were bred and maintained in-house and the offspring were used in this study. After weaning, mice were same sex housed in groups of one to five per cage. Mice were housed in a temperature- (18-23 °C), humidity- (40-60%) and light-controlled (12 light/12 dark) room. Access to food and water was *ad libitum*.

### In vivo electrophysiology

Electrophysiological recordings were performed as previously described (13, 14) with slight modifications. Mice were anesthetized with isoflurane via a precision vaporizer (VetFlo and Kent Scientific) (3.5-4.0% for induction, 0.75-1.25% for maintenance) in 1 L/min oxygen flow. Ophthalmic ointment was placed on both eyes to prevent drying. After loss of reflexes, animals were placed on the recording platform where body temperature was maintained at 36-37 °C through a feedback-controlled heating pad monitored via a rectal thermoprobe (Stoelting). The skull was exposed, and connective tissue was removed to ensure adherence of the headplate. A custom-made Acrylonitrile Butadiene Styrene plastic headplate was tightly secured to the skull using superglue and dental cement. A craniotomy was performed on the right hemisphere, ^~^1.5-2.0 mm lateral from the midline and ^~^1.5 mm anterior from lambda using a branchpoint of the anterior cerebral artery as a reference to target the anteromedial region of SC. Ophthalmic ointment was then fully removed from both eyes. A thin layer of silicone oil was then applied to the eyes to prevent drying while allowing for clear optical transmission.

A 16-channel silicone probe (NeuroNexus Technologies) was lowered between 1.0 and 1.6 mm into the SC at a 45° angle to the horizontal and a 45° angle to the midline. A polytrode electrode with 2 columns of sites (8 sites per column) at 50 μm intervals (model A1-16-Poly2-5mm-50s-177-A16) was used (Figure 1B). The shanks of the probes were 5 mm long, with a maximum width at the top of the shank of 68 μm. Probes were coated in 1,1’-Dioctadecyl-3,3,3’,3’-Tetramethylindocarbocyanine Percholate (DiI, Invitrogen) for post-procedure visualization of probe placement (Figure 1B). Electrical signals were acquired at ^~^25 kHz, amplified and filtered between 0.7-7 kHz using a System 3 workstation (Tucker-Davis Technologies). Signals were analyzed with custom software in MATLAB (MathWorks) and individual units were identified by spike sorting methods using independent components analysis, as previously described (15).

**Fig. 1.**
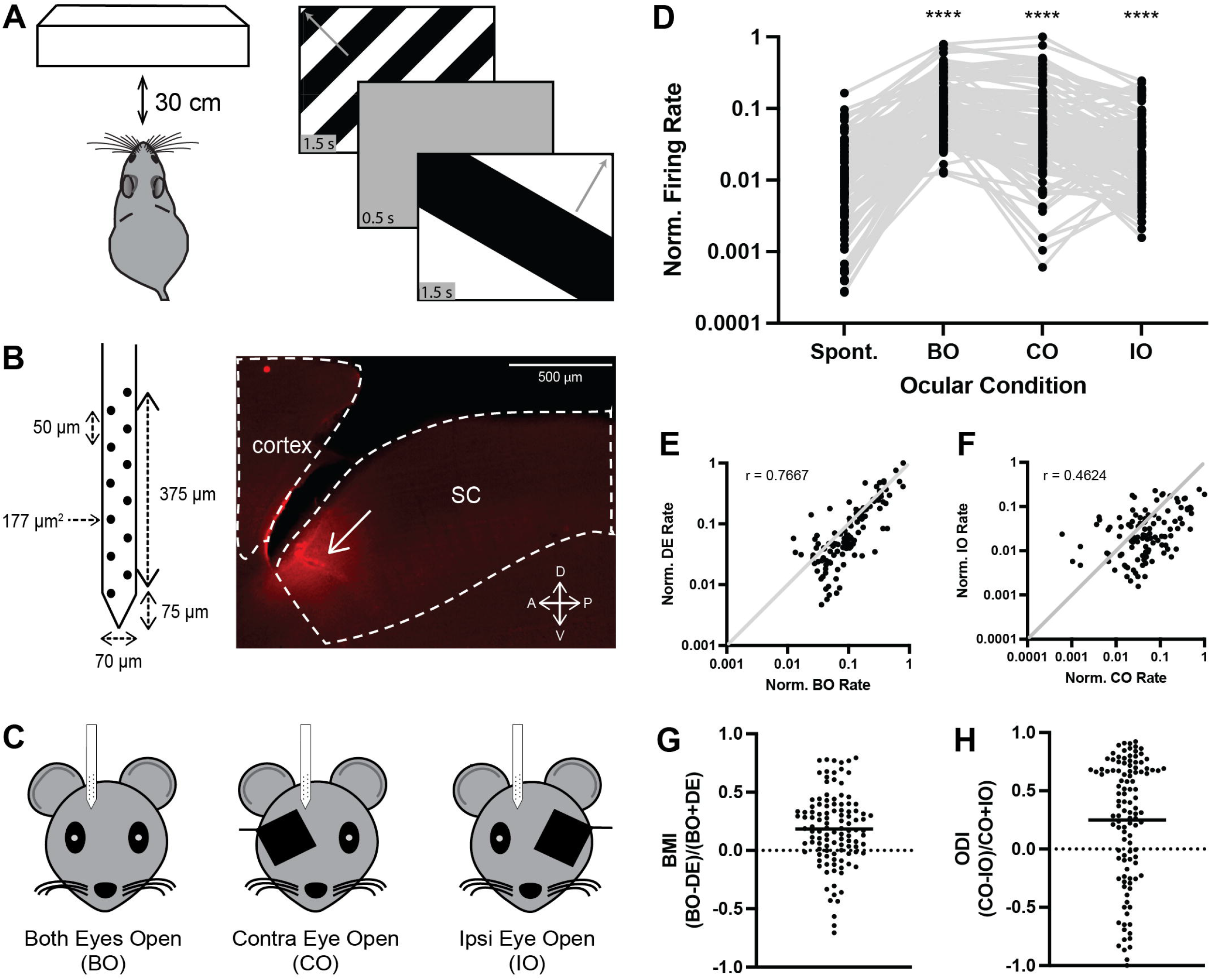
Presentation of visual stimuli in different ocular conditions reveals diverse patterns of neuronal responses. A) Schematic of visual stimulus paradigm. B) Schematic of 16-channel multi-electrode used (*left*) and parasagittal section through the superior colliculus (SC) revealing dye-labeled electrode track in the anterior-medial SC. C) Schematic of ocular conditions in which visual stimuli were presented. D) Quantification of normalized spontaneous and evoked firing rates of all visually responsive units identified (N = 112 units from 25 mice) under each ocular condition. ****, P < 0.0001 vs. spontaneous rate, repeated measures one-way ANOVA with Tukey’s *post hoc* test. E) Comparison of normalized firing rates under the dominant eye (DE) and both eyes open (BO) conditions for each visually responsive unit. F) Comparison of normalized firing rates under the ipsilateral eye open (IO) and contralateral eye open (CO) conditions for each visually responsive unit. G & H) Quantification of the binocular modulation index (BMI) (G) and ocular dominance index (ODI) (H) for all visually responsive units. Solid line represents the mean.

At the end of the recording, the animal was deeply anesthetized and intracardially perfused with ice-cold phosphate buffered saline (PBS) followed by 4% paraformaldehyde in PBS. Brains were sagittally sectioned at 150 μm with a VT1000 vibratome (Leica) and imaged on a BX63 Automatic Fluorescent Microscope (Olympus) for probe placement verification.

### Visual stimuli

Visual stimuli were generated using custom MATLAB software, as previously described (15). The monitor (52 x 29.5 cm, 60 Hz refresh rate, ^~^35-45 cd/m^2^ mean luminance) was placed 30 cm from the eyes, subtending ^~^80° x 50° of visual space and centered on the midline of the animal to maximize coverage of the binocular visual field. (Figure 1A). Visual stimuli consisted of drifting square waves (100% contrast) at 12 different orientations (30° spacing), six different spatial frequencies (SFs) (between 0.01 and 0.32 cycles/deg [cpd] over six logarithmic steps), and one temporal frequency (2 Hz) presented for 1.5 s with an inter-stimulus interval of 0.5 s (Figure 1A). In addition, full-field flash (0 cpd) and blank (gray) screens were presented to provide a robust visual stimulus and determine spontaneous firing rate, respectively. Each stimulus condition was presented in 5-7 trials in a pseudorandom order in three different ocular conditions: 1) binocularly to both eyes (BO), 2) monocularly to the eye contralateral to the recording site (CO), and 3) monocularly to the ipsilateral eye (IO) (Figure 1C). For monocular conditions, an opaque piece of cardboard was placed directly in front of the opposite eye to occlude it from viewing the screen. Following this, animals were again presented visual stimuli in the BO condition to ensure no confounding effect of anesthetic.

### Visual tuning analysis

For each identified unit, we determined if the unit was visually responsive under each ocular condition. To do so, we first determined the preferred stimulus, defined as the stimulus that elicited the highest mean firing rate in each ocular condition. Since we and others have found that spontaneous rates of neurons in the SC are usually low (16), we chose a stringent cutoff to determine if a unit was visually responsive in each ocular condition. A unit was determined to be visually responsive if 1) the response to the preferred stimulus was significantly different from the spontaneous rate based on a one-way ANOVA comparing the spontaneous rate and responses to all orientations shown at the preferred SF (if the preferred stimulus was a drifting grating) or Wilcoxon’s Rank Sum test (if the preferred stimulus was full-field flash), 2) the mean firing rate elicited by the preferred stimulus was greater than 2 standard deviations above the spontaneous rate of the unit, 3) the firing rate elicited by the preferred stimulus was greater than 2 standard deviations above the spontaneous rate in at least two-thirds of the trials, and 4) the firing rate elicited by the preferred stimulus exceeded 5 Hz. Units that were determined to be visually responsive under any ocular condition were utilized for further analysis. Since we presented stimuli in the BO ocular condition twice, we chose the epoch in which the sum of the mean rates across all visual stimuli was highest for further analysis.

For visually responsive units, we calculated the degree of orientation and directional tuning. To quantify the degree of orientation and direction selectivity, the global orientation and direction selectivity indices were calculated (gOSI and gDSI, respectively) (17, 18). gOSI and gDSI were calculated as the vector sum of responses normalized by the scalar sum of responses. gOSI was calculated as: 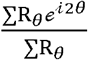, where R_θ_ is the mean firing rate at θ direction of gratings. gDSI was calculated as: 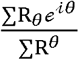. A cell was considered orientation selective (OS) if it had a gOSI greater than 0.2 and a gDSI less than 0.2, while a cell was considered direction selective (DS) if it had a gDSI greater than 0.2. For OS and DS units, we determined the preferred spatial frequency as that which had the highest mean rate across all directions shown and the preferred direction after fitting the tuning curve with a wrapped Gaussian to firing rates at each direction presented at the preferred spatial frequency (15). From this, we determined the preferred orientation for OS units by first normalizing the preferred direction to half of the full phase, i.e. 0° to 180°, then adding or subtracting 90° to determine the orthogonal (16). In addition, we determined the tuning width as the half-width at half-maximum of the fitted curve above the baseline and the F_1_/F_0_ ratio as the firing rate at the stimulus frequency divided by the mean rate.

### k-means Clustering

As input for the clustering algorithm, we utilized the firing rates of individual units under each ocular condition, normalized to that which elicited the highest rate, and two indices that compared interactions between the eyes: the binocular modulation index (BMI) and ocular dominance index (ODI). BMI was calculated as: 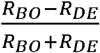, where *R_BO_* is the rate elicited by the preferred stimulus under the BO ocular condition and *R_DE_* is the peak rate elicited by the dominant eye, i.e. the greater of the rates elicited by the preferred stimulus under CO and IO ocular conditions. ODI was calculated as 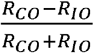, where *R_CO_* and *R_IO_* are the rates elicited by the preferred stimulus under the CO and IO ocular conditions, respectively. Clustering was performed using the ‘kmeans’ function in Matlab, utilizing the default settings for distance measure (squared Euclidean) and centroid seed locations (random data point), which yielded cluster assignments and distances to the centroid for each visually responsive unit. Silhouette coefficients were determined using the ‘silhouette’ function in Matlab, utilizing the default settings for distance (squared Euclidean). We performed five independent iterations of *k*-means clustering with accompanying silhouette analysis

### Statistical Analysis

Statistical analysis was conducted on N=25 male and female animals. There was no difference between male and female responses. Each animal had one to nine visually responsive units, totaling N=112 neurons for analysis. All values are reported as mean ± standard error of the mean (SEM). All data were tested for normal distribution using the Anderson-Darling test. Comparisons of mean values between ocular conditions or putative clusters was performed via a one-way ANOVA and multiple comparisons performed *post hoc* via a Tukey’s test. Correlations were determined by Spearman’s test. Slopes of best fit lines were determined via least squares fit (LSF) and compared to the line of identity via an extra sum of squares F test (ESSF). All analyses and graph plotting were done with Prism (GraphPad Software).

## Results

### Neurons in the anteromedial SC exhibit varied responses to distinct ocular presentations

To determine eye-specific influence on visual responsiveness in the anteromedial SC, we presented drifting square waves of varying orientation, direction of movement, and spatial frequency on a screen placed directly in front of animals while recording multiunit activity from a 16-channel silicone probe placed in the anteromedial SC (Fig. 1A & B). Mice were exposed to visual stimuli in 3 different ocular conditions: 1) both eyes open (BO) 2) only the contralateral eye to the recording site open (CO) or 3) only the ipsilateral eye open (IO) (Fig. 1C). To quantify relative effects of ocular condition, we determined the peak firing rate of each visually responsive elicited by the stimulus in each condition (N = 112 units from 25 mice), as well as the spontaneous firing rate (Fig. 1D). These rates were then normalized to the maximal rate across all units and ocular conditions. We observed a main effect of ocular condition on normalized firing rate (nFR) (*P* < 0.0001, repeated measures one-way ANOVA). Specifically, the BO, CO and IO conditions elicited a significantly higher firing rate than the spontaneous rate (mean nFR ± SEM: spontaneous = 0.0157 ± 0.0022, BO = 0.1487 ± 0.0148, CO = 0.1105 ± 0.0154, IO = 0.0462 ± 0.0051; BO or CO or IO vs. Spont., *P* < 0.0001, Tukey’s multiple comparisons test). Furthermore, peak nFR elicited under the BO condition was significantly greater than that under either the CO and IO conditions (BO vs. CO, *P* = 0.0001; BO vs. IO, *P* < 0.0001, Tukey’s), and the peak CO nFR was greater than the peak IO nFR (*P* < 0.0001, Tukey’s). These differences in elicited responses suggest a potential interaction between eye-specific inputs.

We next compared the firing rates under the dominant eye condition (DE), i.e. the monocular condition (CO or IO) that elicited the higher rate, to that under the BO condition and found that these rates were highly correlated (Spearman r = 0.7667, *P* < 0.0001) (Fig. 1E), suggesting substantial contributions of the DE to responses when stimuli were presented to both eyes. Of note, we observed multiple points in each of these comparisons that fell well above or below the line of identity, indicating possible facilitation or suppression of visual response by the non-dominant eye. We also compared the firing rates elicited under CO and IO monocular conditions and found a lower, but significant, degree of correlation (Spearman r = 0.4624, *P* < 0.0001) (Fig. 1F). Again, many points fell well above the line of identity, suggesting a subset of neurons in the anteromedial SC differentially respond to the contralateral and ipsilateral eye.

To further explore potential variations in firing due to ocular-specific modulation, we calculated two indices to describe the responses under each ocular condition. First, we determined the binocular modulation index (BMI), which compares the BO rate to the DE rate. That is, a neuron predominantly driven by the DE would have a BMI near zero, whereas a neuron potentiated or suppressed under the BO condition would have a BMI near 1 or −1, respectively. Intriguingly, we observed a wide range of BMI values (maximum = 0.7949; minimum = −0.7059; standard deviation [SD] = 0.3131), suggesting neurons in the anteromedial SC can exhibit both positive and negative interactions between the dominant and nondominant eye (Fig. 1G). Interestingly, these data suggest that influence of the non-dominant eye is more often facilitation, since the mean was positive (mean BMI = 0.1838 ± 0.0296), and the 75^th^ percentile value (0.3801) was closer to the maximum than the 25^th^ percentile value (−0.009) was to the minimum.

Next, we determined the ocular dominance index (ODI), which compares the firing rate under each monocular condition. For this metric, a neuron equally driven by the contralateral and ipsilateral eyes would have an ODI near zero, while a neuron predominantly driven by the contralateral or ipsilateral eye would have an ODI near 1 or −1, respectively. Again, we observed a wide range of ODI values (mean = 0.2500 ± 0.0499; maximum = 0.9218; minimum = −0.9500; SD = 0.5266), suggesting neurons could be driven by either eye (Fig. 1H). Most were driven more by the contralateral eye than the ipsilateral eye, as 67.9% of ODI values were positive. Taken together, these data suggest that neurons in the anteromedial SC can exhibit varied responses to visual stimuli presented under different ocular conditions. Furthermore, most are predominantly driven by the contralateral eye, and the non-dominant eye is more likely to be potentiating.

### Identification of five distinct ocularly-modulated response profiles in the SC

We next sought to determine if the varied response profiles to distinct ocular presentations of visual stimuli could be grouped into putative subtypes of binocularly modulated neurons in the anteromedial SC. To begin, we plotted the BMI and ODI of individual units and found broad dispersion in the resulting scatter plot (Fig. 2A). Next, we performed *k*-means clustering based on the BMI and ODI of individual units, along with their unit-specific nFRs. Since the number of clusters identified in *k*-means clustering is determined *a priori*, we tested a range of *k* values from 1 to 7 and performed 5 iterations of clustering for each value.

**Fig. 2.**
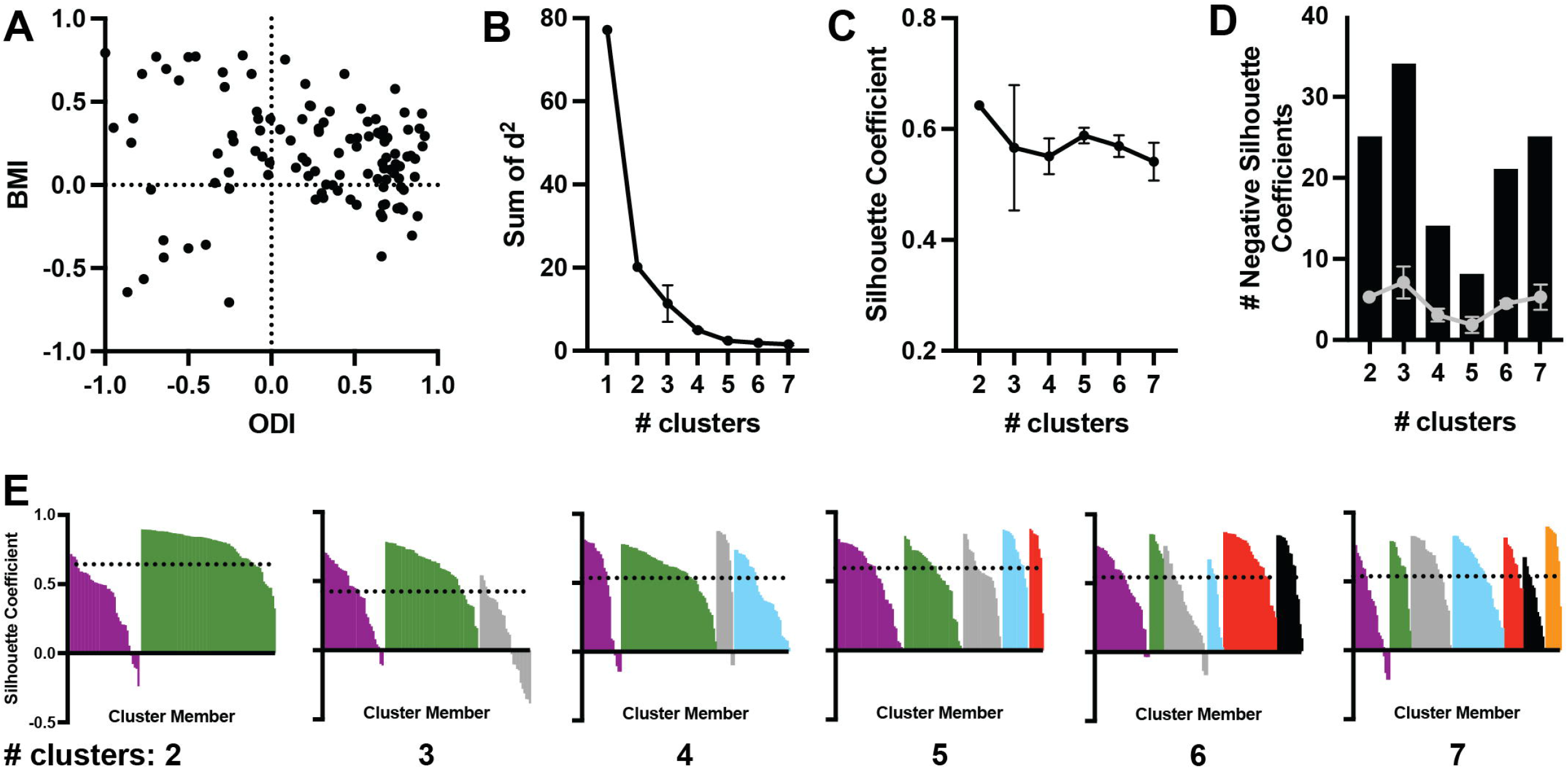
Unbiased clustering of neuronal responses based on differential responses under varied ocular conditions. A) Comparison of binocular modulation index (BMI) and ocular dominance index (ODI) for each visually responsive unit. B-D) Quantification of the sum of squared distances between units and assigned cluster centroid (B), the mean silhouette coefficient across clusters (C), and the cumulative (*black bars*) and mean (*gray dots*) number of negative silhouette values (D) for indicated number of clusters across five iterations of k-means clustering. Data in panels B-D are presented as mean +/- SD. E) Representative plots of individual silhouette coefficients for units in their assigned cluster for the indicated number of clusters.

To evaluate which *k* value best describes the number of clusters in our dataset, we determined the sum of squared distances between individual points and their centroid (Fig. 2B). The resulting plot reveals substantial decreases in value as the number of clusters increases from 1 to 5, but little difference thereafter. In addition, we calculated the mean silhouette coefficient, based on the distances between points within a cluster and the nearest neighboring cluster (Fig. 2C). We found the highest mean silhouette coefficient when *k* = 2, and the second highest when *k* = 5. Since negative silhouette coefficients indicate an increased likelihood that a given point may belong to a different cluster, we also determined the cumulative and mean number of negative silhouette coefficients for 5 iterations of clustering for each value of *k* (Fig. 2D). We observed that the lowest cumulative and mean number of negative silhouette coefficients was found when *k* = 5. To further parse the quality of clustering for different values of *k*, we manually inspected silhouette plots for each value from a representative iteration (Fig. 2E). We observed that when *k* = 2, the vast majority of values above the mean (Fig. 2E, dotted line) belonged to one cluster. Similarly, when *k* = 3, 6 or 7, at least one cluster was comprised of a substantial number of members whose silhouette coefficient was well below the mean (Fig. 2E: Cluster 3.3 [gray], Cluster 6.3 [gray], Cluster 7.1 [purple]). In contrast, when *k* = 4 or 5, all clusters were comprised of a substantial number of members well above or very near the mean (Fig. 2E). Based on the relatively low sum of squared distances, high mean silhouette coefficient, low number of negative silhouette coefficients, and robustness of each cluster relative to the mean silhouette coefficient, we chose a value of *k* = 5 for further analysis.

To interrogate the validity of our choice for number of clusters, we first plotted the BMI as a function of ODI for all units with assigned clusters indicated by distinct colors and numbered 1-5 in descending order of population (Fig. 3A). Broadly, units that had low BMI and low ODI values were grouped together (Cluster 5.5, red), as were units with high values for each (Cluster 5.2, green). A third group with high BMI but low ODI values appeared to be grouped together (Cluster 5.4, cyan). A fourth group with intermediate BMI and ODI values was identified (Cluster 5.3, gray), as well as a fifth group with high ODI but intermediate BMI values (Cluster 5.1, purple). Overall, clusters with high ODI values (5.1 and 5.2) represented a majority of units identified (33.04 and 27.68%, respectively; 60.72%, total), followed by the cluster with intermediate ODI values (5.3, 18.75%) (Fig. 3B). Those clusters with low ODI values (5.4 and 5.5), i.e. driven predominantly by the ipsilateral eye, represented the smallest proportion of units (13.39% and 7.14%, respectively) (Fig. 3B). We next asked if the individual clusters could be differentiated based on their BMI and ODI values. Indeed, we found a main effect of cluster on BMI (*P* < 0.0001, one-way ANOVA), with significant differences determined between each group (mean BMI ± SEM, 5.1: −0.0234 ± 0.0226, 5.2: 0.3748 ± 0.0235, 5.3: 0.1952 ± 0.0294, 5.4: 0.6296 ± 0.0466, 5.5: −0.4316 ± 0.0755; all pairwise comparisons, *P* < 0.0001, Tukey’s) (Fig. 3C). Furthermore, we found a main effect of cluster on ODI (*P* < 0.0001, one-way ANOVA), with significant differences found between those clusters with high, intermediate, and low ODI values (mean ODI ± SEM, 5.1: 0.6512 ± 0.0291, 5.2: 0.5499 ± 0.0421, 5.3: −0.0407 ± 0.0405, 5.4: −0.5739 ± 0.0747, 5.5: −0.6002 ± 0.0720; 5.1 or 5.2 vs. 5.3 or 5.4 or 5.5, *P* < 0.0001; 5.3 vs. 5.4 or 5.5, *P* < 0.0001; Tukey’s) (Fig. 3D). Taken together with our analyses of *k* values, these data support the possibility that five distinct putative subtypes of ocularly modulated neurons may be identified in the anteromedial SC.

**Fig. 3.**
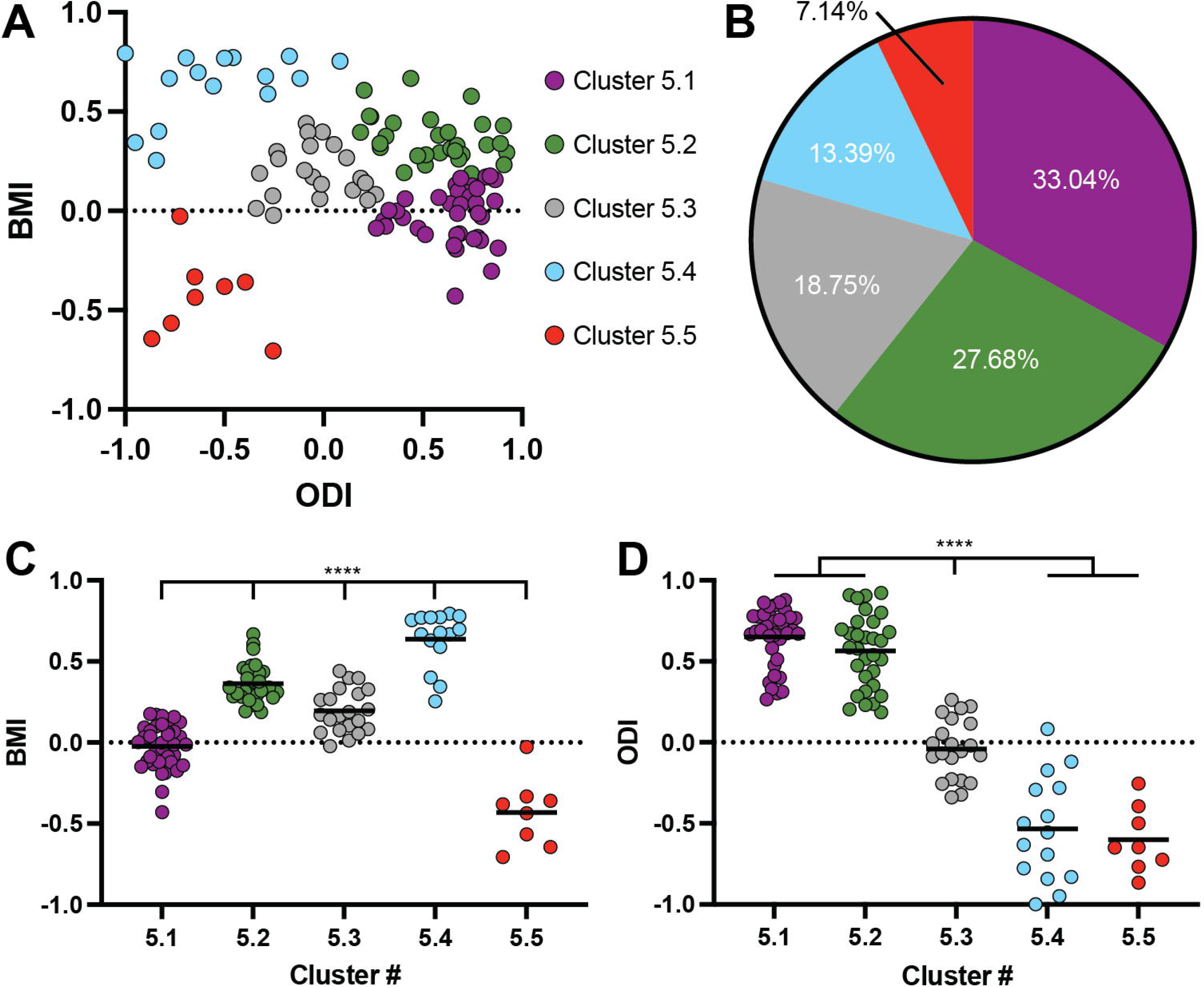
Proportions of putative cell types based on differential responses to visual stimuli presented in varied ocular conditions. A) Comparison of binocular modulation index (BMI) and ocular dominance index (ODI) for each visually responsive unit with putative clusters indicated by color. B) Proportion of indicated clusters amongst all visually responsive units. C & D) Quantification of BMI (C) and ODI (D) for each cluster. Solid lines indicate mean. ****, P < 0.0001 between indicated groups, one-way ANOVA with Tukey’s *post hoc* test.

### Putative subtypes exhibit distinct patterns of response to visual stimuli presented in different ocular conditions

To further evaluate the composition of putative ocularly modulated subtypes in the SC, we examined the responses of the units in each cluster to visual stimuli presented under different ocular conditions. Analysis of peri-stimulus spike time histograms and raster plots suggested varied responses for each cluster to different ocular conditions (Fig. 4A-E). Indeed, for all clusters, we found a main effect of ocular condition on nFR (5.1, *P* < 0.0001; 5.2, *P* < 0.0001; 5.3, *P* < 0.0001; 5.4, *P* = 0.002; 5.5, *P* = 0.0008; repeated measures one-way ANOVA) (Fig. 4A’-E’). However, subsequent analysis of the differences in normalized firing rates under spontaneous, BO, CO, and IO conditions for each cluster revealed distinct profiles of activation (Fig. 4A’-E’).

**Fig. 4.**
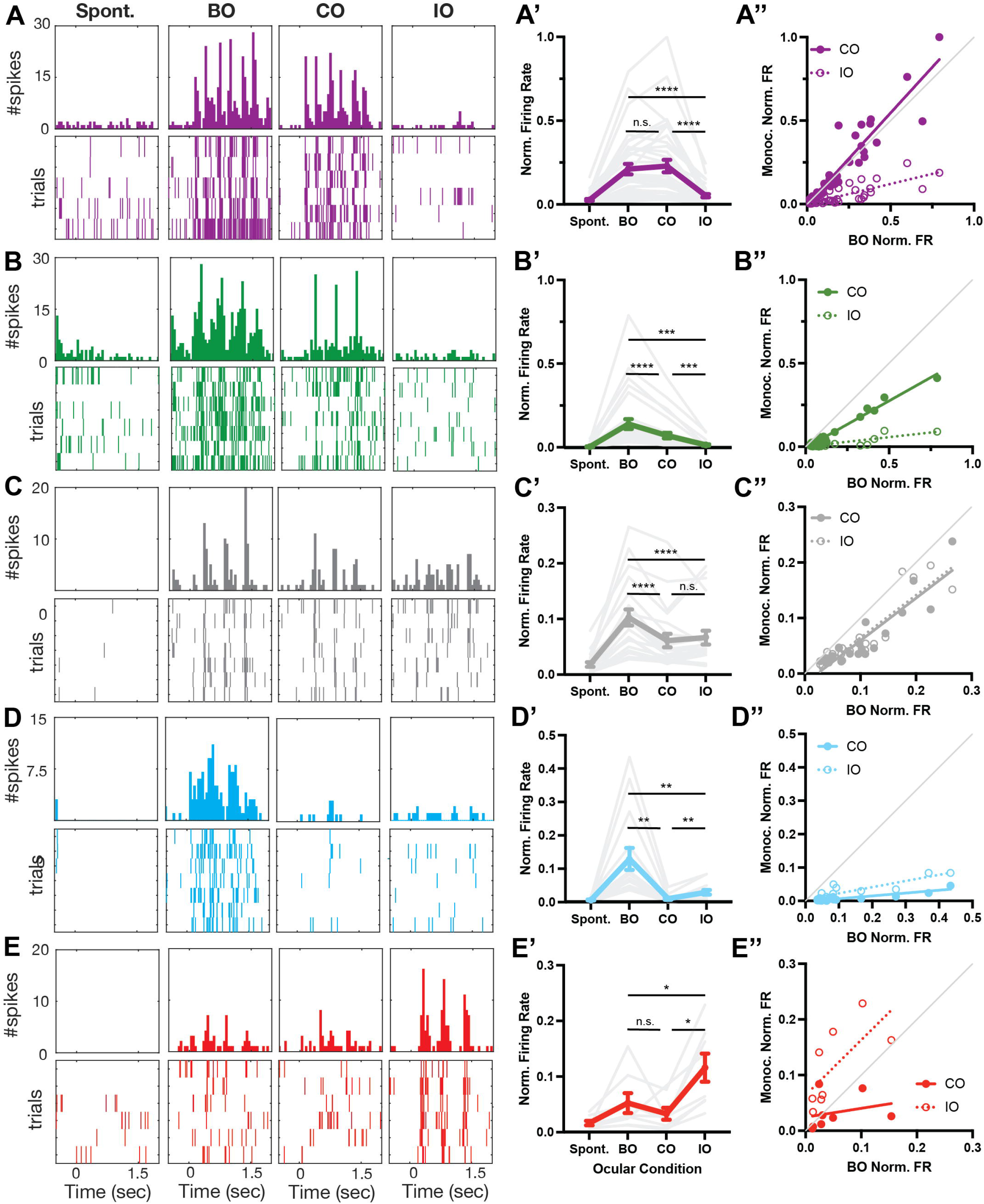
Distinct response profiles under varied ocular conditions of putative cell types. **A-E)** Peri-stimulus spike time histograms (*top*) and raster plots of individual trials (*bottom*) of representative neuronal responses to a blank screen (Spont.) and the most effective visual stimulus in the indicated ocular condition (BO, both eyes open; CO, contralateral eye open; IO, ipsilateral eye open). A’-E’) Plots of the mean normalized firing rate (*thick, colored lines*) for each cluster under the indicated ocular condition. Responses of individual units are depicted by thin, gray lines. *, p < 0.05; **, p < 0.01; ***, p < 0.001; ****, p < 0.0001; n.s. no significant difference between indicated groups, one-way ANOVA with Tukey’s *post hoc* test. A’’-E’’) Comparison of monocular firing rate (*solid dots, CO*; *open circles, IO*) with the firing rate in the BO condition.

For a first subgroup of clusters, including Clusters 5.1, 5.2, and 5.3, the peak nFR elicited under all ocular conditions was significantly greater than the spontaneous rate (mean nFR ± SEM, 5.1-Spont.: 0.0257 ± 0.0056, 5.1-BO: 0.2093 ± 0.0312, 5.1-CO: 0.2274 ± 0.0369, 5.1-IO = 0.0509 ± 0.0095; vs. 5.1-Spont., 5.1-BO: *P* < 0.0001, 5.1-CO: *P* < 0.0001, 5.1-IO: *P* = 0.0004; 5.2: Spont.: 0.007 ± 0.0019; BO, 5.2-CO: 0.0716 ± 0.0165, 5.2-IO: 0.0175 ± 0.0039; vs. 5.2-Spont., 5.2-BO: *P* = 0.0002, 5.2-CO: *P* = 0.0005, 5.2-IO: *P* = 0.001; 5.3-Spont.: 0.0183 ± 0.0041, 5.3-BO: 0.1026 ± 0.0145, 5.3-CO: 0.0612 ± 0.0122, 5.3-IO: 0.0666 ± 0.0123; vs. 5.3-Spont., 5.3-BO: *P* < 0.0001, 5.3-CO: *P* = 0.0007, 5.3-IO: *P* = 0.0003; Tukey’s). However, these clusters could be distinguished based on the differences in elicited rates. For Cluster 5.1, we found that the peak BO and CO rates were not different from one another (*P* = 0.2058, Tukey’s), but both were significantly greater than the peak IO rate (BO or CO vs. IO, *P* < 0.0001, Tukey’s) (Fig. 4A’). In contrast, for Cluster 5.2, we found that the peak BO rate was significantly great than both the CO and IO rates (BO vs. CO: *P* < 0.0001, BO vs. IO: *P* = 0.0002, Tukey’s), and the CO rate was greater than the IO rate (*P* = 0.0007, Tukey’s) (Fig 4B’). Similarly, the peak BO rate of Cluster 5.3 was significantly greater than both the peak CO and IO rates (BO vs. CO: *P* < 0.0001, BO vs. IO: *P* < 0.0001, Tukey’s), but the CO and IO rates were not different from one another (*P* = 0.4713).

For a second subgroup of clusters, including Clusters 5.4 and 5.5, not all elicited rates were significantly different from the spontaneous rate. For Cluster 5.4, firing rate was significantly higher that the spontaneous rate under the BO and IO conditions, but not the CO condition (mean nFR ± SEM, 5.4-Spont.: 0.0053 ± 0.0016; 5.4-BO: 0.1335 ± 0.0347; 5.4-CO: 0.0091 ± 0.0033; 5.4-IO: 0.0296 ± 0.0072; vs. 5.4-Spont., 5.4-BO: *P* = 0.0088, 5.4-CO: *P* = 0.1986, 5.4-IO: *P* = 0.0088, Tukey’s). Further, the rates elicited under BO and IO conditions were significantly higher than that observed under the CO condition (BO vs. CO: *P* = 0.0088, IO vs. CO: *P* = 0.0063, Tukey’s). For Cluster 5.5, neither the BO nor CO rate was significantly greater than spontaneous, but the IO rate was (mean nFR ± SEM, 5.5-Spont.: 0.0159 ± 0.0041; 5.5-BO: 0.0518 ± 0.0178; 5.5-CO: 0.0326 ± 0.0108; 5.5-IO: 0.1154 ± 0.0253; vs. 5.5-Spont., 5.5-BO: *P* = 0.2086, 5.5-CO: *P* = 0.2427, 5.5-IO: *P* = 0.0179, Tukey’s). The peak IO rate was also greater than that elicited under both the BO and CO conditions (IO vs. BO: *P* = 0.0385, IO vs. CO: *P* = 0.0188, Tukey’s).

To further discern potential differences in responses in putative ocularly modulated subtypes, we compared the peak monocular (CO and IO) rates to the peak binocular (BO) rate (Fig. 4A’’-E’’). For Cluster 5.1, we found that both the CO and IO rates were correlated with the BO rate (CO, Spearman r = 0.9623, *P* < 0.0001; IO, Spearman r = 0.8026, *P* < 0.0001) Interestingly, we found that the slope of a best fit line through the CO-BO comparison was not significantly different from the line of identity (slope =1.096, least squares fit [LSF]; *P* = 0.2032, Extra sum-of-squares F test [ESSF]), whereas that for the IO-BO comparison was (slope = 0.2418, LSF; *P* < 0.0001, ESSF), suggesting that firing under the BO condition for units in Cluster 5.1 is predominantly driven by the contralateral eye, while the ipsilateral eye minimally influences firing.

For Cluster 5.2, again both monocular rates were correlated with the BO rate (CO, Spearman r = 0.8781, *P* < 0.0001; IO, Spearman r = 0.6536, *P* < 0.0001). In contrast to Cluster 5.1, the slopes of best fit lines through both the CO-BO and IO-BO comparison for Cluster 5.2 were significantly different from the line of identity (CO-BO: slope = 0.5633, *P* < 0.0001; IO-BO: slope = 0.1110, *P* < 0.0001; LSF, ESSF). The relatively high slope of the CO-BO fit suggests that evoked firing under the BO condition may be largely driven by the contralateral eye, but the deviation from the line of identity suggests it may be potentiated by the ipsilateral eye.

We found that the monocular rates for Cluster 5.3 were also highly correlated with the induced BO rate (CO, Spearman r = 0.8746, *P* < 0.0001; IO, Spearman r = 0.9015, *P* < 0.0001). Intriguingly, while the slopes of the best fit line for each were close to 1, they were significantly different from the line of identity (CO-BO, slope = 0.7646, *P* = 0.0096; IO-BO, slope = 0.7665, *P* =0.0135; LSF, ESSF). These data suggest that both the contralateral and ipsilateral eye contribute to firing under the BO condition for units in Cluster 5.3, but neither is more dominant than the other.

For Cluster 5.4, we again found that firing in both the CO and IO conditions were significantly correlated with the BO rate (CO, Spearman r = 0.7137, *P* = 0.0054; IO, Spearman r = 0.8801, *P* < 0.0001); however, the slopes of both best fit lines were low and significantly different from the line of identity (CO-BO, slope = 0.08679, *P* < 0.0001; IO-BO, slope = 0.1815, *P* < 0.0001; LSF, ESSF). Since stimulation under the IO condition elicits a small but significant increase in firing rate, these data suggest that the ipsilateral eye may contribute slightly more to the response observed under BO conditions, but that it is substantially potentiated by stimulation of the contralateral eye.

Finally, for Cluster 5.5, we found that the IO rate, but not the CO rate, was well correlated with the BO firing rate (CO, Spearman r = 0.5238, *P* = 0.1966; IO, Spearman r = 0.7619, *P* = 0.0368). Consistent with the possibility that inputs from the ipsilateral eye drive a substantial portion of firing of units in Cluster 5.5, the slope of a best fit line through the comparison with BO rate was not significantly different from the line of identity (slope = 0.9821, LSF; *P* = 0.9719, ESSF). Further, the slope of the CO-BO comparison for Cluster 5.5 was significantly different from the line of identity (slope = 0.1597, LSF; *P* = 0.0126, ESSF). Together, these data suggest that units in Cluster 5.5 may be predominantly driven by ipsilateral inputs, and that contralateral inputs may be suppressive.

### Orientation- and direction-selectivity of putative ocularly modulated subtypes in the SC

We next explored the tuning properties of putative subtypes of neurons in the anteromedial SC. We chose to elicit visual responses under different ocular conditions using a drifting square wave stimulus, which is a strong driver of visual responses in the SC and allowed us to determine the selectivity of units for orientation or direction of movement. To assess orientation tuning, we calculated the global orientation selectivity index (gOSI), and found a main effect of cluster identity on gOSI (*P* = 0.0239, one-way ANOVA) (Fig. 5A). Subsequent analysis revealed the gOSI of Cluster 5.2 was significantly greater than that of Cluster 5.5 (mean gOSI ± SEM, 5.2: 0.3338 ± 0.0200, 5.5 = 0.1837 ± 0.0588, *P* = 0.0394, Tukey’s *post hoc* test), but no other pairwise differences were observed. These data also suggest that many units in Clusters 5.1, 5.3 and 5.4 are orientation-selective (OS), since the mean gOSI for each is greater than our cut-off for selectivity of 0.2 (5.1: 0.2564 ± 0.0205, 5.3: 0.2835 ± 0.0371, 5.4: 0.3174 ± 0.0260).

**Fig. 5.**
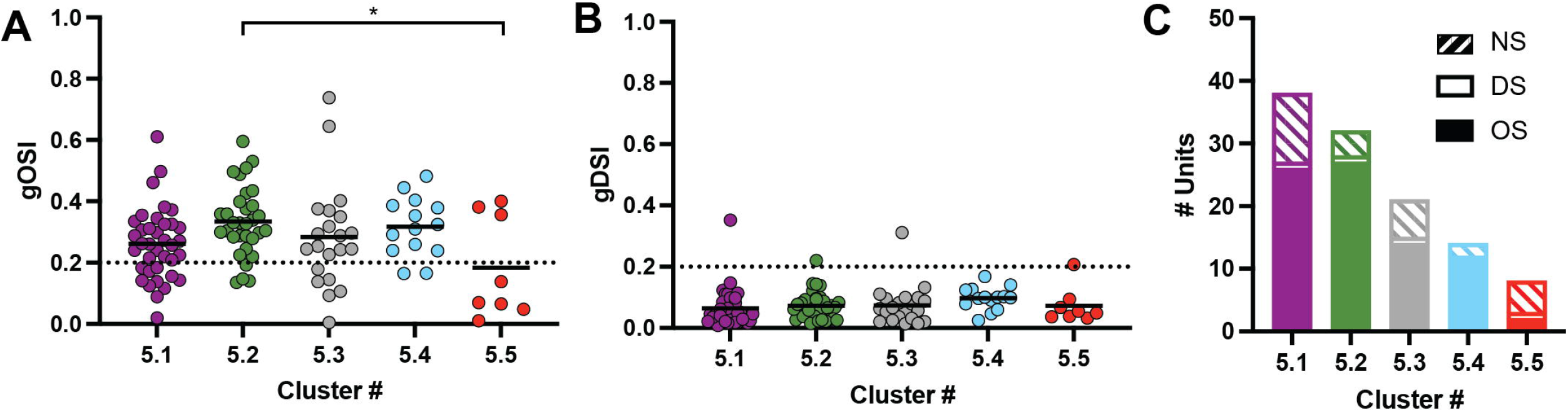
Orientation- and direction-selectivity of putative ocularly modulated cell types in the SC. A & B) Quantification of the global orientation and direction selectivity indices (gOSI, A; gDSI, B). Solid line represents the mean, dotted line at 0.2 represents cut-off for selectivity. *, P < 0.05 between indicated groups, one-way ANOVA with Tukey’s *post hoc* test. C) Cumulative numbers of non-selective (NS, *striped bars*), orientation-selective (OS, *solid bars*), and direction-selective (DS, *open bars*) units identified in each putative cell type.

In addition to orientation tuning, we evaluated selectivity for motion by calculating the global direction selectivity index (gDSI) (Fig. 5B). In contrast to orientation-selectivity, we did not find a main effect of cluster identity for gDSI (*P* = 0.4412, one-way ANOVA). In fact, none of the clusters exhibited robust direction-selectivity, since the mean gDSI for each was well below our cutoff of 0.2 (mean gDSI ± SEM, 5.1: 0.0634 ± 0.0100; 5.2: 0.0722 ± 0.0080; 5.3: 0.0739 ± 0.0141; 5.4: 0.0968 ± 0.0100; 5.5: 0.0721 ± 0.0205). Overall, we identified only four direction-selective (DS) units, defined as those with a gDSI > 0.2, and no cluster had more than one DS unit (Fig. 5C). In contrast, we found a high proportion of units in 4/5 clusters were OS, defined as a gOSI > 0.2 and gDSI < 0.2 (5.1: 25/37, 67.6%; 5.2: 27/32, 84.4%; 5.3:14/21, 66.7%; 5.4: 12/14, 85.7%; 5.5: 2/8: 25.0%) (Fig. 5C). Taken together, these data suggest that a substantial proportion of neurons in the anteromedial SC are selective for orientation, regardless of their differential patterns of ocular modulation.

### Comparison of tuning properties of OS units in putative ocularly modulated subtypes

To discern if the OS units in each cluster could be distinguished from one another, we next interrogated their tuning properties specifically. We excluded Cluster 5.5 from these analyses, since only two units met our cut-off for OS. First, we determined the preferred orientation of each OS unit by fitting a 2-D Gaussian to the tuning curve and adjusting values such that they fell in a 0-180° range, where 0/180° represented a horizontal grating and 90° represented a vertical one. Consistent with previous studies (19), we found that most OS units in the anteromedial SC were tuned to orientations that were rotated counterclockwise from a vertical line (mean preferred orientation ± SEM, 5.1:140.8 ± 4.628°; 5.2:130.6 ± 5.513°; 5.3: 130.2 ± 5.551°; 5.4:140.4 ± 6.073°) (Fig. 6A). Indeed, we found no effect of cluster identity on preferred orientation (*P* = 0.3704, one-way ANOVA), suggesting that preferred orientation is not altered based on ocular modulation properties.

**Fig. 6.**
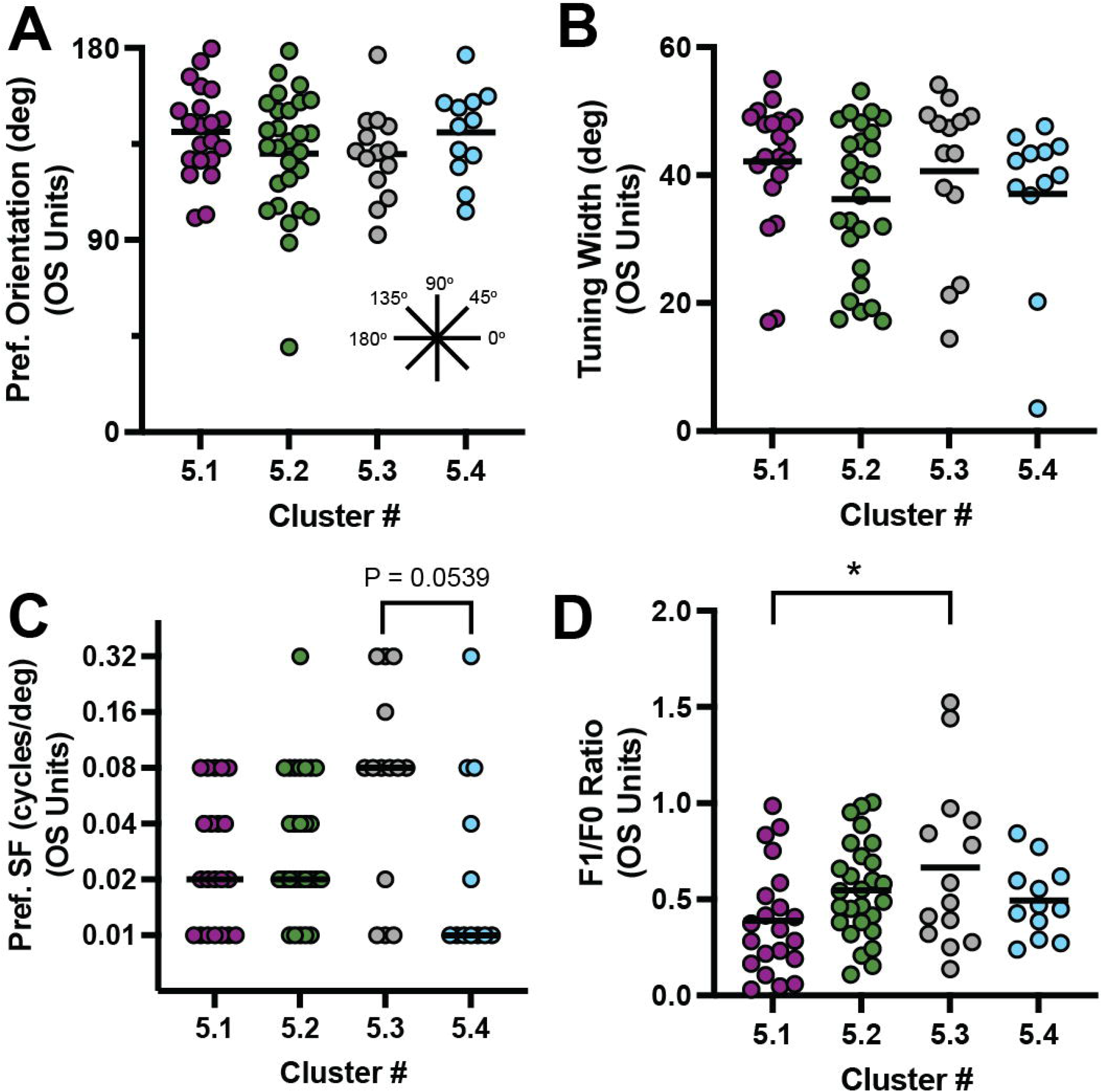
Similar tuning properties of OS cells across putative ocularly modulated cell types. **A-D)** Quantification of the preferred orientation of drifting square wave stimulus (A), tuning width (B), preferred spatial frequency (SF) (C), and linearity (F1/F0 ratio) for orientation-selective (OS) units in each of the indicated clusters. For panels A, B, & D, solid line represents the mean. *, P < 0.05, one-way ANOVA with Tukey’s *post hoc* test. For panel C, solid lines represent the median. Cumulative distributions were analyzed via a Kruskal-Wallis test, which indicated a main effect of cluster identity (P = 0.0434), and Dunn’s multiple comparison test was used *post hoc*.

Next, we examined the sharpness of tuning of OS units towards their preferred direction by calculating the tuning width of a best fit curve (Figure 6B). In general, OS neurons of all putative subtypes were sharply tuned (mean tuning width ± SEM, 5.1:42.11 ± 2.202°; 5.2: 36.24 ± 2.228°; 5.3: 40.63 ± 3.345°; 5.4: 37.06 ± 3.668°). And, we found no effect of cluster identity on tuning width (*P* = 0.3079, one-way ANOVA), suggesting that differences in ocular modulation are not related to sharpness of tuning.

We next explored the spatial frequency (SF) preferences of OS units in each cluster (Fig. 6C). We defined the preferred SF as that which elicited the highest mean firing rate across all orientations shown. We observed that most OS units exhibited a preference for lower spatial frequencies, save for those in Cluster 5.3 (median preferred SF, 5.1: 0.02; 5.2: 0.02; 5.3: 0.08; 5.4: 0.01). Indeed, we found a main effect of cluster identity on the cumulative distribution of preferred SFs (*P* = 0.0434, Kruskal-Wallis test), and subsequent analysis revealed a trend towards a shift in distribution towards higher SFs for OS units in Cluster 5.3 compared to those in Cluster 5.4 (*P* = 0.0539, Dunn’s multiple comparisons test). These data suggest that OS units in Cluster 5.3 may be tuned to higher SFs that those in other clusters.

Finally, we explored the linearity of OS units in each putative subtype by determining the ratio of firing at the first harmonic (F_1_) to the mean rate (F_0_) (20). Consistent with previous data (13, 16), we observed that most OS units in the anteromedial SC were non-linear, having F_1_/F_0_ ratios below one (mean F_1_/F_0_ ratio ± SEM, 5.1: 0.3889 ± 0.0614; 5.2: 0.5477 ± 0.0481; 5.3: 0.6666 ± 0.1156; 5.4: 0.4933 ± 0.0552) (Fig. 6D). Intriguingly, we found a trend towards an effect of cluster identity on F_1_/F_0_ ratio (*P* = 0.0546, one-way ANOVA). Indeed, subsequent analysis revealed that the mean F_1_/F_0_ ratio for OS units in Cluster 5.3 was significantly greater than those in Cluster 5.1 (*P* = 0.0383, Tukey’s *post hoc* test). These data suggest that OS units in Cluster 5.3 may respond more linearly to visual stimuli than those in other clusters.

### Relationships of ocular-specific orientation tuning reflect ocular modulation of visual response

Thus far, we have shown that neurons in the anteromedial SC exhibit varied patterns of response to visual stimuli presented in different ocular conditions. Since many of these units are OS, we next wanted to explore whether the orientation tuning of units under different ocular conditions might provide insight into the mode of ocular modulation. To do so, we evaluated the ocular condition-specific gOSIs of all units, as well as the preferred orientations of OS units, in each cluster (Fig. 7). Similar to our analyses of firing rates (Fig. 4), we observed distinct patterns of correlation for each measure between clusters.

**Fig. 7.**
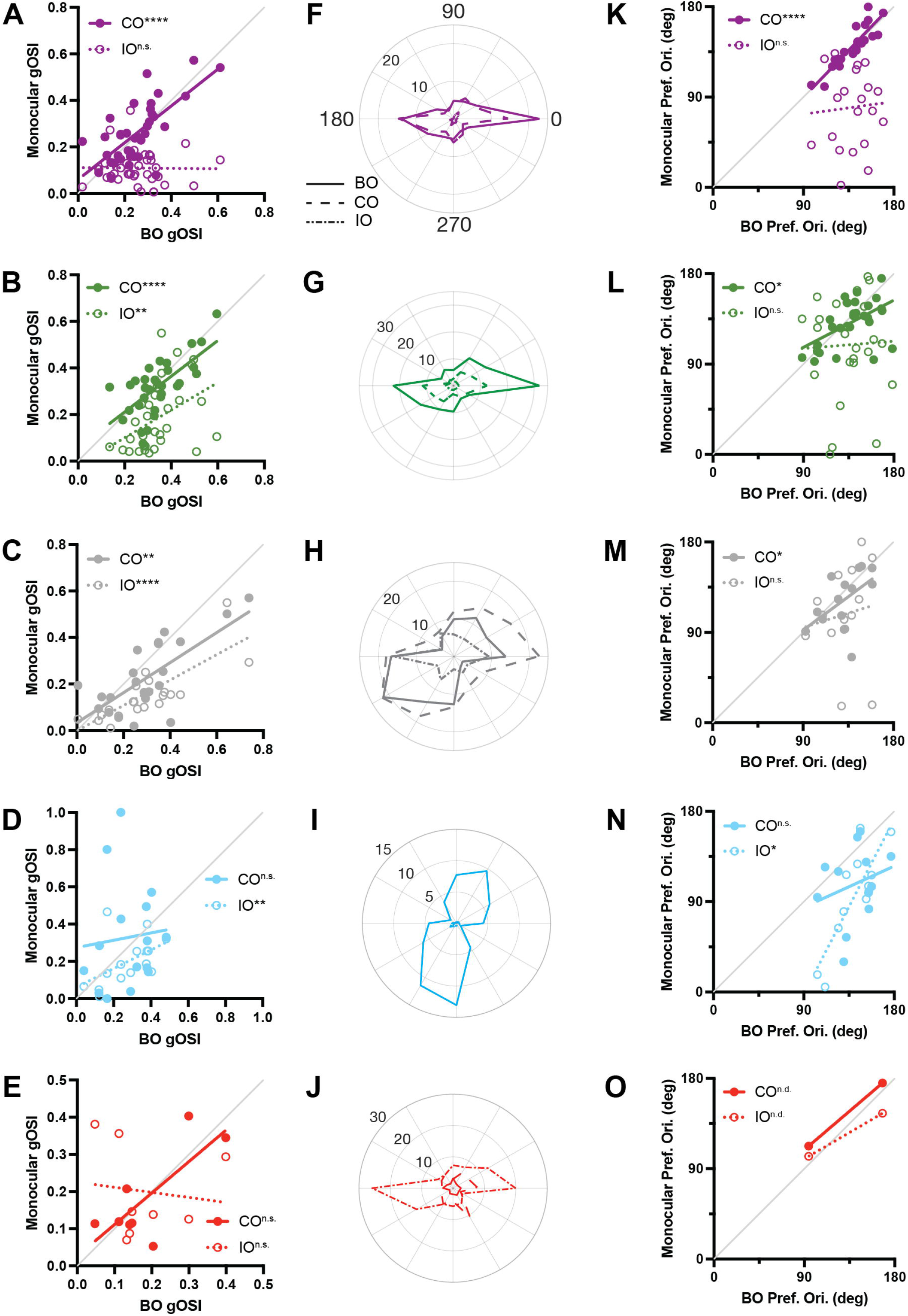
Putative ocularly modulated cell types exhibit distinct patterns of correlation between monocular and binocular orientation tuning properties. **A-E)** Comparison of global orientation selectivity index (gOSI) under monocular (*solid dots, CO*; *open circles, IO*) visual stimulation and gOSI under binocular visual stimulation. **, P < 0.01; ****, P < 0.0001; n.s., not significant correlation between monocular and binocular gOSI, Spearman’s correlation test. F-J) Polar plots of representative OS units from each putative cell type in which the response under both eyes open (BO, *solid lines*), contralateral eye open (CO, *dashed lines*), and ipsilateral eye open (IO, *stippled lines*) conditions are overlaid. K-O) Comparison of monocular (*solid dots, CO*; *open circles, IO*) and binocular preferred orientation for orientation-selective units in each cluster. *, P < 0.05, Spearman’s correlation test.

For Cluster 5.1, we found that the gOSI under CO conditions was highly correlated with the BO gOSI (Spearman r = 0.6399, *P* < 0.0001), but that the gOSI under the IO condition was not (Spearman r = −0.0638, *P* = 0.7077) (Fig. 7A). Furthermore, we found that the slope of the best fit line through the CO:BO gOSI comparison was not significantly different from the line of identity (slope = 0.79, LSF; *P* = 0.0882, ESSF), while that of the IO:BO comparison was (slope = - 0.008, LSF; *P* < 0.0001, ESSF). Similarly, when we compared the tuning curves of OS units in Cluster 5.1 under each ocular condition, we observed a striking similarity between the BO and CO conditions (Fig. 7F). Indeed, we found a highly significant correlation between preferred orientation in the BO and CO conditions (Spearman r = 0.9195, *P* < 0.0001). As these units exhibited minimal evoked response under the IO condition, neither the tuning curve nor preferred orientation showed similarity to those observed under the BO condition (Spearman r = −0.0013, *P* = 0.9955). Taken together with correlations observed in peak evoked firing rates, these data suggest that the visual response and tuning of units in Cluster 5.1 under BO conditions is predominantly driven by the contralateral eye.

For Cluster 5.2, we found that the gOSI under both CO and IO conditions was correlated with the BO gOSI (CO: Spearman r = 0.6484, *P* < 0.0001; IO: Spearman r = 0.5000, *P* = 0.0042) (Fig. 7B). Interestingly, the best fit line through either dataset was not significantly different from the line of identity, though that for the IO:BO comparison trended towards difference (CO:BO: slope = 0.7675, *P* = 0.0991; IO:BO: slope = 0.5940, *P* = 0.0593; LSF, ESSF). Consistent with this, the tuning curves under BO and CO conditions of OS units in Cluster 5.2 appeared similar, while that observed under the IO condition was generally weak (Fig. 7G). Indeed, we found that the preferred orientation of OS units in Cluster 5.2 under the CO condition was significantly correlated with that under the BO condition (Spearman r = 0.4719, *P* = 0.0129), while that under the IO condition was not (Spearman r = 0.0586, *P* = 0.7715) (Fig. 7L). However, despite the fact that the slope of a best fit line through the CO:BO comparison of preferred orientations was high, it was significantly different from the line of identity (slope = 0.5221, LSF; *P* = 0.0109, ESSF). Taken together with relationships between evoked firing, these data suggest that visual responses and tuning of units in Cluster 5.2 under BO conditions is largely, but not completely, driven by inputs through the contralateral eye.

For Cluster 5.3, the gOSI under both CO and IO conditions was highly correlated with the BO gOSI (CO: Spearman r = 0.5753, *P* = 0.0064; IO: Spearman r = 0.7922, *P* < 0.0001) (Fig. 7C). However, the slope of the best fit line through both the CO:BO and the IO:BO datasets, despite being high, were significantly different from the line of identity (CO:BO: slope = 0.6455, *P* = 0.0056; IO:BO: slope = 0.5447, *P* < 0.0001; LSF, ESSF). We observed that the tuning curves for most OS units in Cluster 5.3 were similar under all ocular conditions (Fig. 7H). Indeed, the preferred orientation observed under CO conditions was significantly correlated with that found under BO conditions (Spearman r = 0.6000, *P* = 0.0261) and the slope of a best fit line was not different from the line of identity (slope = 0.7416, LSF; *P* = 0.4747, ESSF) (Fig. 7M). While the preferred orientation under IO conditions was not significantly correlated with the BO condition (Spearman r = 0.3758, *P* = 0.1862), the slope of a best fit line through the data points was not significantly different from the line of identity (slope = 0.3173, LSF; *P* = 0.3457, ESSF). Taken together with correlations between evoked firing rates, these data suggest that inputs from both the contralateral and ipsilateral eye contribute to the response and tuning of units in Cluster 5.3 under BO conditions.

For Cluster 5.4, the gOSI under the IO condition was significantly correlated with the BO gOSI, but that found under the CO condition was not (CO: Spearman r = 0.2357, *P* = 0.3966; IO: Spearman r = 0.6679, *P* = 0.008) (Fig. 7D). However, the slope of the best fit line through the CO:BO comparison was not significantly different from the line of identity, while that of the IO:BO comparison was (CO-BO: slope = 0.1973, *P* = 0.1136; IO-BO: slope = 0.4987, *P* = 0.0143; LSF, ESSF). Intriguingly, manual inspection of tuning curves under each ocular condition revealed that most OS units in Cluster 5.4 exhibited a discernable tuning curve under the BO condition (Fig. 7I). Despite this, we found that the preferred orientation of OS units in Cluster 5.4 under IO conditions was significantly correlated with that elicited under BO conditions (Spearman r = 0.6573, *P* = 0.0238), while that found under CO conditions was not (Spearman r = 0.2168, *P* = 0.4990) (Fig. 7N). However, the best fit line through the IO:BO comparison was significantly different from the line of identity (slope = 1.931, LSF; *P* = 0.0458, ESSF). Taken together with correlations between evoked firing under all ocular conditions, these data suggest that ipsilateral, but not contralateral, inputs contribute to the response and tuning of units in Cluster 5.4. However, the tuning may be shaped by non-dominant contralateral inputs.

Finally, for Cluster 5.5, we found that the gOSI determined under neither the CO nor IO conditions were significantly correlated with the BO gOSI (CO: Spearman r = 0.3571, *P* = 0.3894; IO: Spearman r = −0.2619, *P* = 0.5364) (Fig. 7E). Intriguingly, a best fit line through the BO:CO comparison was not significantly different than the line of identity, while a line through the IO:CO comparison was (CO:BO: slope = 0.8479, *P* = 0.6233; IO:BO: slope = −0.1362, *P* = 0.0431; LSF, ESSF). Since only two units in Cluster 5.5 met our OS criteria, further evaluation of correlation between preferred orientation in different ocular conditions was not possible (Fig. 7O). Of note, manual inspection of the tuning curves of these units revealed a robust response under IO conditions, but minimal activation in other contexts (Fig. 7J). Taken together with correlations between evoked firing rates, these data support the possibility that visual responses of units in Cluster 5.5 are primarily driven by the ipsilateral and that those from the contralateral eye, though suppressive, may help shape orientation tuning.

## Discussion

The mouse SC is a midbrain structure that receives visual information from the vast majority of RGCs, including inputs from both the contralateral and ipsilateral eye. However, the ways in which eye-specific inputs modulate visual responses and tuning properties of neurons in the SC are not well understood. Here, we characterized the responses of visual neurons in the anteromedial SC to stimuli presented monocularly and binocularly. In contrast to previous studies, we found that a substantial portion of neurons exhibited interactions between eye-specific inputs, as well as evidence of prominent activation by ipsilateral stimulation. Clustering based on indices reflecting binocular modulation and ocular dominance revealed 5 distinct profiles of eye-specific response modulation. Four of the five identified groups exhibited robust selectivity for orientation of stimulus. Of these, the OS units in one group (Cluster 5.3) preferred higher SFs and responded more linearly than those in other groups. Comparison of orientation tuning between monocular and binocular stimulus presentations confirmed the distinct profiles of ocular modulation and provide insights into potential mechanisms underlying these phenomena. Together, these data suggest that binocular interactions in the mouse SC are more common and varied than previously assumed.

### Prevalence of BN neurons in the mouse SC

Previous studies of visual tuning properties in the rodent SC suggested minimal binocular interactions or even ipsilaterally-influenced responses (6, 7). In contrast, we found a surprising proportion of the units we identified in the anteromedial mouse SC exhibited binocular modulation, manifested as both facilitation and suppression by the non-dominant *eye*. Furthermore, while most units observed were more driven by stimulation of the contralateral eye (61.61%, Clusters 5.1 and 5.2), we found substantial numbers of units for which the ipsilateral eye was dominant or equally contributing to visual responses (38.39%, Clusters 5.3, 5.4 and 5.5). The lack of evidence for binocularity in the rodent SC in previous studies could be driven by experimental methods or analysis strategies or a combination of both.

First, while the mouse SC receives bilateral input from both contralateral and ipsilateral RGCs, the contralateral RGC input to the SC is dense while the ipsilateral input is sparse. Previous studies have shown that contralateral terminals assume a volume of ^~^401×10^-6^ mm^3^ while ipsilateral terminals only assume 9.83×10^-6^ mm^3^ (21). And, these inputs are segregated into distinct laminae in the SC. Eye-specific segregation itself does not preclude binocularity, as has been demonstrated in the dorsal lateral geniculate nucleus (dLGN) (22–24). But, the organization of ipsilateral terminals in the mouse dLGN may lend itself to easier identification of binocular neurons. In the mouse, ipsilateral RGC terminals are clustered in the center of the bean-shaped dLGN and surrounded on all sides by contralateral RGCs. Combined with the convoluted, interwoven nature of dendritic and axonal terminals (25), this organization provides ample opportunity for convergence of inputs from both eyes. Indeed, recent trans-synaptic retrograde tracing data suggests this is possible (26). In contrast, the mouse SC is a much larger structure and ipsilateral RGCs are spread along a crescent spanning the anterior and medial borders. Furthermore, ipsilateral RGC terminals are organized into clusters, rather than a contiguous lamina, further reducing potential interactions. Thus, the probability of penetrations in early studies targeting these regions specifically may have been reduced, especially if the anteromedial regions was not intentionally sampled.

A second contributing factor could be derived from the classification strategy we utilized to determine binocular interactions. Previous studies in which abundant binocular interactions were reported in the anterior SC of hamsters relied upon visual stimuli eliciting a response when presented through each eye (8). While we did find a subpopulation of neurons in the mouse SC, namely Cluster 5.3, that exhibited responses in both CO and IO conditions, the majority had more complex responses to presentation in multiple ocular conditions. For instance, units in Clusters 5.1 and 5.2 both exhibited robust responses to stimulation of the contralateral eye, but minimal responses to stimulation of the ipsilateral eye. However, we observed a significant difference in BMI between these groups, providing a mechanism to distinguish these populations. It is likely that these units would have been presumed to belong to a common subtype of contralaterally-driven neurons in previous studies. Of note, the responses we observed were elicited in the anesthetized state. Given the robust differences in visual tuning in the awake state (27), it will be interesting to determine if binocular interactions in the SC are consistent in different contexts.

### Varied modes of ocular modulation in the anteromedial SC

Perhaps even more surprising than the observation that ipsilateral stimulation may be more robust in the SC than previously presumed is the fact that we observed distinct modes of binocular interactions. These variations raise the intriguing possibility that visual neurons in the SC may be comprised of multiple ocularly-modulated subtypes and that the neuronal wiring to produce such varied responses may be even more complex. To begin, it is important to note that the most numerous subpopulation we encountered, Cluster 5.1 (33.03% of units), did not exhibit any binocular interactions. That is, the evoked firing rates for these units were strikingly similar when stimuli were presented binocularly or monocularly to the contralateral, as were the tuning to drifting square waves and the preferred orientation of OS units. These data suggest that the most common pattern of wiring in the anteromedial SC involves neurons that are exclusively innervated by RGCs from the contralateral eye that provide excitatory input.

The second most numerous subpopulation we encountered, Cluster 5.2 (28.57% of units), was also driven to a great degree by contralateral input. However, presentation of stimuli binocularly revealed a facilitation, which is supported by the small, but significant, increase in evoked firing when stimuli were presented to the ipsilateral eye only. Such a pattern suggests that neurons in Cluster 5.2 may receive dense innervation from contralateral RGCs, as well as sparse innervation from ipsilateral ones, each providing excitatory input. Unfortunately, previous studies in which binocular units were encountered in the rodent SC did not explore the phenomenon of facilitation, reporting only that stimulation of the ipsilateral eye elicited “weak” influence (8). However, investigations of binocular interactions in the cat SC revealed that facilitation was common by the non-dominant eye (28), supporting the possibility that this type of response can be achieved.

Next, we encountered a subpopulation of units, Cluster 5.3, that exhibited nearly equal response to both contralateral and ipsilateral stimulation, which appeared to be additive when stimuli were presented binocularly. Such a response suggests neurons in this cluster may receive equally dense innervation from each eye, both of which are excitatory in nature. Given that this type of profile would have been prototypical for a binocularly-driven unit in classical studies (6–8), it is surprising that more were not encountered. Indeed, nearly 20% of all visually responsive units reported here exhibited this profile. We did observe that OS units in Cluster 5.3 tended to prefer smaller spatial frequencies and were more linear in response, so one potential cause for not encountering them could be the use of suboptimal stimuli.

In addition to units responding strongly to contralateral inputs as observed in Clusters 5.1, 5.2 and 5.3, we found two other subpopulations that exhibited unexpected response profiles. The first, Cluster 5.4, had a weak response to ipsilateral stimulation (and none to contralateral) but was robustly activated when stimuli were presented binocularly. Such a response is reminiscent of “obligatory binocular” units previously reported in secondary visual cortex of primates (29–31), which coded for binocularity disparity, allowing for stereopsis. While we did not explore disparity selectivity here, recent work suggests that the SC may be critical for a prey capture behavior requiring binocular vision (10, 11). Indeed, we recently reported that a mouse model lacking ipsilateral RGC input to the SC showed similar disruptions in prey capture efficiency (12). Taken together, these data raise the intriguing possibility that neurons in the SC displaying binocular-dependent or -modulated responses may be critical for complex behavioral responses.

Finally, we identified a cluster of cells that exhibited robust responses to stimulation of the ipsilateral eye, but not under other conditions. While we encountered only a handful of these cells, there segregation from other clusters was very evident. In contrast to others, units in Cluster 5.5 were the only to exhibit suppression by the non-dominant eye, suggesting an inhibitory element to the wiring underlying these responses. One potential source for inhibition could be GABAergic RGCs, a subset of which project to the SC (32, 33). Alternatively, multiple visual cortical areas project topographically to the SC and in alignment with the retinocollicular map (34), providing one potential source of binocular input. However, studies have shown that binocular integration in the dLGN is independent of cortical feedback (23, 35). Intriguingly, binocular responses are prevalent in the frog tectum, which arise not from direct ipsilateral RGC inputs, but rather indirectly from the nucleus isthmi (36). The mammalian homologue of the nucleus isthmi, the parabigeminal nucleus, also projects to the SC and could convey binocular information. Future studies leveraging eye- or region-specific manipulations of activity are needed to resolve these possible circuit mechanisms.

### Conclusion

Here, we report the extent of binocular interactions in the mouse SC. Contrary to previous reports, we observed multiple distinct modes of binocular interactions. In the majority of units, a strong response to contralateral eye stimulation was observed, however, we found that the non-dominant ipsilateral eye could provide additive or facilitative influence under binocular stimulation conditions. Furthermore, smaller subpopulations that exhibited either robust facilitation or suppression by the non-dominant contralateral eye were identified. Taken together, these data reveal a more prevalent and complex array of binocular interactions in the mouse SC than previously thought, opening the door for understanding how these subpopulations may contribute to binocular- and SC-dependent behaviors.

